# Dynamic species classification of microorganisms across time, abiotic and biotic environments — a sliding window approach

**DOI:** 10.1101/105395

**Authors:** Frank Pennekamp, Jason I. Griffiths, Emanuel A. Fronhofer, Aurélie Garnier, Mathew Seymour, Florian Altermatt, Owen L. Petchey

## Abstract

1. Technological advances have greatly simplified to take and analyze digital images and videos, and ecologists increasingly use these techniques for trait, behavioral and taxonomic analyses. The development of techniques to automate biological measurements from the environment opens up new possibilities to infer species numbers, observe presence/absence patterns and recognize individuals based on audio-visual information.

2. Streams of quantitative data, such as temporal species abundances, are processed by machine learning (ML) algorithms into meaningful information. Machine learning approaches learn to distinguish classes (e.g., species) from observed quantitative features (phenotypes), and in-turn predict the distinguished classes in subsequent observations. However, in biological systems, the environment changes, often driving phenotypic changes in behaviour and morphology.

3. Here we describe a framework for classifying species under dynamic biotic and abiotic conditions using a novel sliding window approach. We train a random forest classifier on subsets of the data, covering restricted temporal, biotic and abiotic ranges (i.e. windows). We test our approach by applying the classification framework to experimental microbial communities where results were validated against manual classification. Individuals from one to six ciliate species were monitored over hundreds of generations in dozens of different species combinations and over a temperature gradient. We describe the steps of our classification pipeline and systematically explore the effects of the abiotic and biotic environments as well as temporal effects on classification success.

4. Differences in biotic and abiotic conditions caused simplistic classification approaches to be unsuccessful. In contrast, the sliding window approach allowed classification to be highly successful, because phenotypic differences driven by environmental change could be captured in the learning algorithm. Importantly, automatic classification showed comparable success compared to manual identifications.

5. Our framework allows for reliable classification even in dynamic environmental contexts, and may help to improve long-term monitoring of species from environmental samples. It therefore has application in disciplines with automatic enumeration and phenotyping of organisms such as eco-toxicology, ecology and evolutionary ecology, and broad-scale environmental monitoring.

## 1 Introduction

Society is presently in the midst of an automation revolution that was initiated in the middle of the 20th century by the invention of the Turing machine. Tasks once performed by humans are steadily being relinquished to computers that are more efficient at tedious jobs than their human counterparts. Likewise, ecologists are increasingly relying on semi- or fully automated monitoring systems to collect images, videos and sounds to characterize environments and biological interactions. The field of animal biometrics develops quantitative approaches to describe and identify species and individuals, using morphological traits and behaviours from audio-visual sources (Kühl and Burghardt 2013). Examples include species monitoring using audio (Russo and Voigt 2016, Depraetere et al. 2012) or visual information (Weinstein 2015, Swinnen et al. 2014), identification based on patterns such as color or shape (Karanth et al. 2006), or behaviour from movement trajectories and associated accelerator data (Nathan et al. 2012). Whereas these approaches show promise for cataloging different aspects of biodiversity (MacLeod et al. 2010), they require careful optimization to accurately measure species abundance and phenotypic variation (Russo and Voigt 2016).

Digital image and video analysis comprises a set of techniques to perform time intensive tasks, including counting, measuring and tracking individuals (Pennekamp and Schtickzelle 2013, Dell et al. 2014). Given the ongoing development of such automation analyses, image analysis is primarily used under controlled laboratory conditions, whereby populations and individuals are phenotyped (e.g., using movement patterns) to conduct a variety of ecological and evolutionary experiments (Pennekamp and Schtickzelle 2013, Mallard et al. 2013). Nevertheless, in natural systems, image and video based techniques have applied to identify plankton species in marine surveys (Bell and Hopcroft 2008, Culverhouse et al. 2006) and to monitor microorganisms in waste water treatment plants (Amaral et al. 2008; 2004). The wealth of data produced by image and video analysis is both a blessing and a curse. Processing and analysis of the data can become overwhelming when hundreds or thousands of videos are processed and when manual steps are needed as to optimize or supplement the automated work flow (Kühl and Burghardt 2013).

Regardless of whether images, videos or sounds are used, making information available to researchers requires transforming the raw data (e.g., pixel intensity, movement trajectories or frequency and length of calls) into biologically meaningful information (e.g., number of species observed, the individuals present in a specific area, or behavioural patterns). This transformation can be achieved by machine learning techniques such as classification or regression (Tarca et al. 2007, Peters et al. 2014). Machine learning algorithms use quantitative properties such as the pixel intensity, or features of the objects identified by the image analysis step (e.g., size or shape) to predict the class of an object (e.g., to which species an individual belongs). Supervised learning algorithms are trained on data whose class is known (i.e., labeled) and the goal is to accurately predict unknown (i.e., unlabeled) observations. An important prerequisite for training classifiers is hence that the training data adequately describes the properties of the unknown data.

Populations and communities often show considerable interspecific variation in abundance and intraspecific variation in phenotypic traits (Ozgul et al. 2009), both of which may impair reliable species level identification. Phenotypic variation is influenced by intraspecific response to the abiotic and biotic environment, which may induce phenotypic changes in other species within a given community (McGill et al. 2006). Predation, for instance, can alter prey size distributions (Travis et al. 2014, Blumenshine et al. 2000), induce the development of defensive traits (Agrawal 2001), or induce changes in movement strategies (e.g., emigration, diapause) (Preisser et al. 2005). Changes in phenotypic expression may also occur as a response to the abiotic environment as well as species interactions occurring at the same trophic level (Agrawal 2001). Consequently, visual species identification methods need to account for dynamic changes in phenotypes to provide accurate biological classifications.

Microcosms are widely used experimental systems to assess ecological and evolutionary influences on temporal and spatial population and community dynamics (Altermatt et al. 2015, Jessup et al. 2004, Benton et al. 2007), and have been instrumental in testing ecological theory (Cadotte et al. 2005, Altermatt et al. 2015). They have been widely used to test the effects of inter- and intraspecific interactions (Jiang and Kulczycki 2004, Leary and Petchey 2009) and phenotypic plasticity (Pennekamp et al. 2014, Hammill et al. 2010).

Here we develop, apply and validate a novel framework to automate species classification which can account for shifting phenotypic traits in response to environmental change. We examine whether accounting for trait differences of individuals from differing biotic and abiotic environments can permit individuals to be correctly categorized as species in a reasonably complex community. Our approach considers the dynamic nature of the classification, which complements previous attempts that focused on classification of ciliates in simpler communities without environmental variation (Pennekamp et al. 2015, Soleymani et al. 2015).

## 2 Materials and methods

### 2.1 Experimental set up

We used microcosms with six bacterivorous ciliate species (i.e., *Colpidium striatum*, *Dexiostoma campylum*, *Loxocephalus* sp., *Paramecium caudatum*, *Spirostomum teres*, and *Tetrahymena thermophila*). The six species were cultured in standard protist medium, along with a common freshwater bacterium, *Serratia fonticola*, as a food source. The medium consisted of protist pellets (Carolina Biological Supplies, Burlington, NC) at a concentration of 0.55 gL^-1^ of Chalkley’s medium, and two wheat seeds for slow nutrient release (Altermatt et al. 2015).

Two weeks prior to the start of the experiment, we established fresh protist cultures for each of the six species. A sample of 10mL of stock protist culture was added to 1000mL of fresh protist medium in previously autoclaved 1000mL glass bottles (GL 45, Schott Duran, Germany). Populations were checked to ensure carrying capacity was reached prior to the start of the experiment. During the experiment, communities were kept in 250mL glasses bottles.

Single species replicates (species richness 1) were started at a density of three individuals mL^-1^ in 100mL volume. Multispecies communities were initiated by first making 40mL of medium from stock cultures. For a species richness of 2, this was made up of 20mL of each species; for three species there was 13.3mL of each species, and so on, up to 6.66mL of each species in six species treatments. This was topped up to 100mL by addition of 60mL bacteria inoculated protist medium. The starting densities hence were standardized to a fixed fraction of the species specific carrying capacity, which differed across richness levels (see table S1). Controls (species richness equaling 0) contained 100mL of protist medium with *Serratia fonticola*. Experimental units were randomized for each temperature treatment and placed in climate controlled incubators (Pol-Eko Aparatura, Wodzislaw, Poland).

### 2.2 Experimental design

We designed a randomized blocking experiment with 720 experimental microcosms (i.e., 250mL Duran® bottles) whereby we assessed the effect of ecological complexity (i.e., species richness) and temperature on the abundance and automated identification of six ciliate species. Ecological complexity included seven levels of ciliate species richness (0-6), where 0 level species richness was used as a control group. Since the total number of possible species combinations exceeded the number of feasible experimental units, we randomly selected species combinations for the 3, 4 and 5 species richness levels. The same species richness combinations were repeated in each of the three temperature blocks. We replicated each level of species richness and composition twice for all levels including an additional replication for the two lowest and the highest levels of complexity (Table 1) resulting in 120 experimental units per temperature (15 °C, 17 °C, 19 °C, 21 °C, 23 °C and 25 °C).

### 2.3 Video sampling, particle tracking and processing

Sampling of each experimental unit occurred every day for the first 7 days, then 3 times per week for the following 50 days and a final sampling 7 days later. Sampling took place in two parallel blocks such that half of the experimental units (360 units, 3 temperature blocks) were sampled in sequence on consecutive days. For each sampling event, culture medium was gently agitated, and a subsample of 700 µl was taken, mounted onto a glass slide and covered with a glass lid. The height of the sampling chamber was 600 µm and the area filmed 68.7 mm^2^ resulting in a sampled volume of 41.2 µL. Five second videos (at 25 frames per second) were taken using a stereomicroscope (Leica M205 C) with a 16 × magnification mounted with a digital CMOS camera (Hamamatsu Orca C11440, Hamamatsu Photonics, Japan).

**Table 1.**
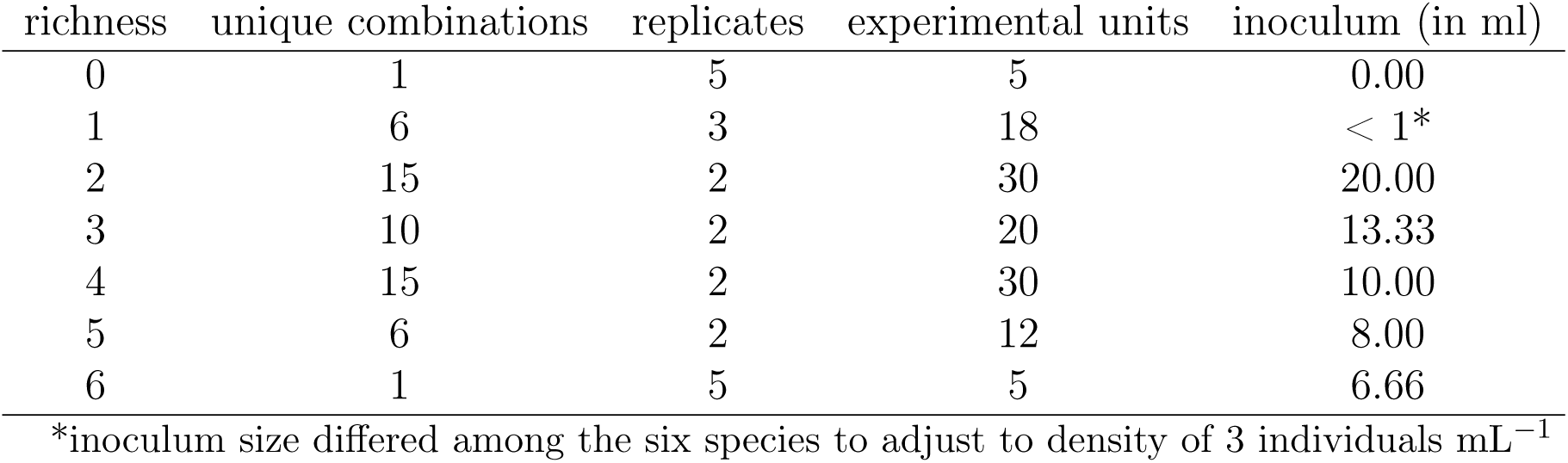
Overview of the experimental design: richness levels, number of unique species combinations per richness level, number of replicates, total of experimental units and inoculum size to start treatments.

| richness | unique combinations | replicates | experimental units | inoculum (in ml) |
| --- | --- | --- | --- | --- |
| 0 | 1 | 5 | 5 | 0.00 |
| 1 | 6 | 3 | 18 | < 1* |
| 2 | 15 | 2 | 30 | 20.00 |
| 3 | 10 | 2 | 20 | 13.33 |
| 4 | 15 | 2 | 30 | 10.00 |
| 5 | 6 | 2 | 12 | 8.00 |
| 6 | 1 | 5 | 5 | 6.66 |

\*inoculum size differed among the six species to adjust to density of 3 individuals mL<sup>-1</sup>

We used the BEMOVI package (version 1.0.2) and the statistical computing environment R (R Development Core Team 2016) to process the 18 720 videos collected during the experiment and extract the raw trajectories (Pennekamp et al. 2015). Global segmentation and tracking parameters were defined for automated processing of videos. The difference lag was defined to be 2s, particle size was restricted to 20 µm to 8100 µm (corresponding to an input of 5 to 2000 pixel in the BEMOVI locate_and_measure_particle function), and the intensity threshold was set to 10. For particle linking, we specified a link range of 0.12 s (3 frames) and a displacement of 81 µm (20 pixels). These settings were optimized using a subset of videos (spanning sampling dates of all single, two and six species combinations at 15 °C, 21 °C and 25 °C). Video settings were optimized to err on the side of including false positives rather than exclude true positives at this step, with exclusion of false positives later in the processing pipeline. For further details regarding video processing, please refer to Pennekamp et al. (2015).

After tracking, trajectories were filtered to remove artifacts such as spurious trajectories (e.g., floating debris). Trajectories for analysis were required to show a minimum net displacement of at least 50 µm, a duration greater than 0.2 seconds, a detection rate of 80 percent (for a trajectory with a duration of 10 frames the individual has to be detected on at least 8 frames), a median step length greater than 2 µm and a minimum mean speed of 50 pixels per second.

### 2.4 Automated species classification in multi-species communities across temperature environments

In supervised classification, data with known classes (i.e., training data, single species cultures in our study) is used to train the classifier (Peters et al. 2014). Training means that the classification algorithm “learns” how to distinguish among known classes based on some quantitative feature such as body size. After training, the classifier can be used to predict the classes of unknown data (i.e., test data, multi-species communities in our study). Reliable training data is hence crucial to construct the classifier. Variation in the training data should only represent biologically meaningful variation, whereas random variation, due to uncontrolled factors, should be minimized. Whereas in small scale experiments, a small number of videos can be individually inspected, in large scale experiments, this is not feasible due large numbers of videos produced. Therefore, our automated species classification pipeline consists of multiple steps (Fig 1). The first three steps were applied in the same way to all videos and can be considered as the data filtering and quality assurance phase. The last three steps are the application of the classification procedure. We apply this procedure using a range of different settings to determine the sensitivity of our results to choices made when performing the machine learning.

**Figure 1.**
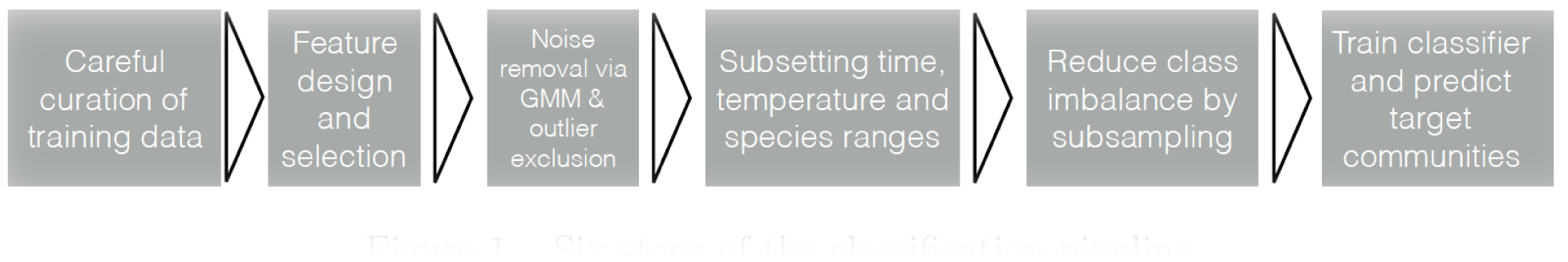
Six steps of the classification pipeline.

#### 2.4.1 Careful curation of training data

During the video recording, microcosms were carefully checked manually for cross-contamination among treatments (e.g., ciliates present in controls, cross-contamination of *Paramecium* monoculture with *Tetrahymena* etc.), and suspect microcosms checked with additional samples. Ten of the 720 of microcosms were contaminated and excluded from further analysis.

A major problem in the analysis of videos is the detection of background (e.g., debris) caused by flow of the sample liquid during videoing. As movement is used to identify the foreground (i.e., the ciliates), spurious observations due to moving background will contribute to incorrect training data. To account for this problem, we implemented a combination of automatic and manual cleaning procedures for the training data. First, we applied more stringent selection for training data by restricting automated trajectory classification to particles moving at speeds greater than 200 µm s^−1^ as moving debris is usually moving at low speed. For classification, all individuals moving faster than 50 µm s^-1^ were considered. Second, we plotted the mean trait values of area and aspect ratio through time to detect outliers and checked these videos manually. After reviewing suspect videos, we excluded inappropriate data. We also used scatterplots of the mean size to define boundaries for each species accounting for the change in morphology through time (see table S2).

#### 2.4.2 Feature selection and feature pre-processing

Features are the quantitative descriptions of the raw trajectories that can be used to distinguish between the different classes by the classifier (Sommer and Gerlich 2013). A large number of features could be calculated from the morphology and trajectory data with some being potentially informative (e.g., size and speed) and others being non-informative (e.g., trajectory length and direction of movement). We identified 10 potential classification features based on our knowledge of characteristics that allow us to discriminate between species. Our 10 classification features were further aggregated at the trajectory level (mean, standard deviation, minimum or maximum), resulting in 15 features (table 2). These 15 features were a subset of features used in previous classification efforts (Pennekamp et al. 2015) and do not include those based on advanced movement features which require substantially longer trajectories for calculation (Soleymani et al. 2015).

All selected variables were checked to have non-zero variance and no missing data. We scaled all features to have zero mean and unit standard deviation, and performed Box-Cox transformation to normalize the data (Kuhn and Johnson 2013). Transformations were applied to the training data for a given community rather than applying the transformations to all individuals at once. Principal component analysis (PCA) was then use to reduce the number of predictor variables under consideration, by obtaining uncorrelated principal components (Quinn and Keough 2002).

#### 2.4.3 Identification of noise and artifacts with Gaussian mixture models (GMMs) and exclusion of background noise

Although the initial filtering removed many data points which reflect background noise, some spurious trajectories remain in the training data and are mixed with the ciliate trajectories. To identify these, we used the control cultures, which were recorded, but did not contain any ciliates and hence only contained spurious trajectories (e.g., due to moving background or changes in light conditions). We trained a Gaussian mixture model (GMM) on the trajectories in the target culture, and identified clusters of spurious trajectories by comparison with the known spurious trajectories from the control cultures.

Depending on the classification algorithm chosen, outliers can have potentially detrimental effects. Given the amount of data available for training the classifier, we decided to only include observations in our training data that fit into the 90% confidence ellipse of a bivariate normal distribution fitted to the first two principal components.

**Table 2.** Morphological and movement features selected for use in classification.

| Code | Type of variable | Measurement method (all are calculated across each of the frames in the particle's trajectory) |
| --- | --- | --- |
| mean_area | Size of particle | Mean area of particle across trajectory |
| sd_area | Temporal variability in particle size | Standard deviation of particle area |
| mean_perimeter | Size and shape of particle | Mean length of perimeter of particle |
| sd_perimeter | Temporal variability of size and shape | Standard deviation of particle perimeter length |
| mean_major | Length of particle | Mean length of major axes of ellipse fitted to particle |
| sd_major | Temporal variability in length | Standard deviation of length of major axis |
| mean_minor | Width of particle | Mean length of minor axis of ellipse fitted to particle |
| sd_minor | Temporal variability in width | Standard deviation of length of minor axis |
| mean_ar | Shape of particle | Mean aspect ratio of particle |
| sd_ar | Temporal variability in shape | Standard deviation of particle aspect ratio |
| sd_turning | Temporal variability of the direction of movement | Circular standard deviation of particle direction |
| gross_speed | Particle speed. | Mean of distance travelled between frames |
| sd_gross_speed | Temporal variability in particle speed. | Standard deviation of distance travelled between frames |
| max_gross_speed | Maximum particle speed. | Maximum distance travelled between frames |
| min_gross_speed | Minimum particle speed. | Minimum distance travelled between frames |

#### 2.4.4 Sub-setting species, date and temperature range for training using a sliding window

Due to environmental change, phenotypes change over time and between environments. This creates a dynamic classification context in which individual features of each category vary spatially and temporally. First, we compared models differing in the number of species used for training. We fitted a model, which contained all the species used in the experiment and an additional noise class that represents the spurious trajectories from the controls, yielding a maximum of seven different classes. We also built a customized model only containing the species we expect in a community plus the noise class, based on our knowledge of the experimental design. Second, we only selected training data within a certain distance in time and temperature (via a sliding window) of the community to be classified. We compared different window sizes (10, 30 and 60 days, i.e. 17%, 50% and 100% of the sampling time) and the temperature range (train based on the temperature of target community vs. all available temperatures). Fig 2 summarizes and illustrates the sliding window approach.

#### 2.4.5 Reduction of imbalance by randomly selecting observations for training

The different ciliate species used in the experiment show variation in cell densities and growth dynamics (Altermatt et al. 2015), hence the number of individuals (i.e., species abundance) differed among species. Imbalance in the number of observations for different classes in the training data can severely decrease the performance of the classifier for the rare class (e.g., low abundance species) (Sommer and Gerlich 2013). Various techniques were developed to deal with this important problem (Kotsiantis et al. 2006). We used a random under-sampling scheme to the majority class to achieve balanced numbers of observations for all classes in the training data. We compared the effect of considering either 250 or 1000 randomly selected observations per species on the classification success.

**Figure 2.**
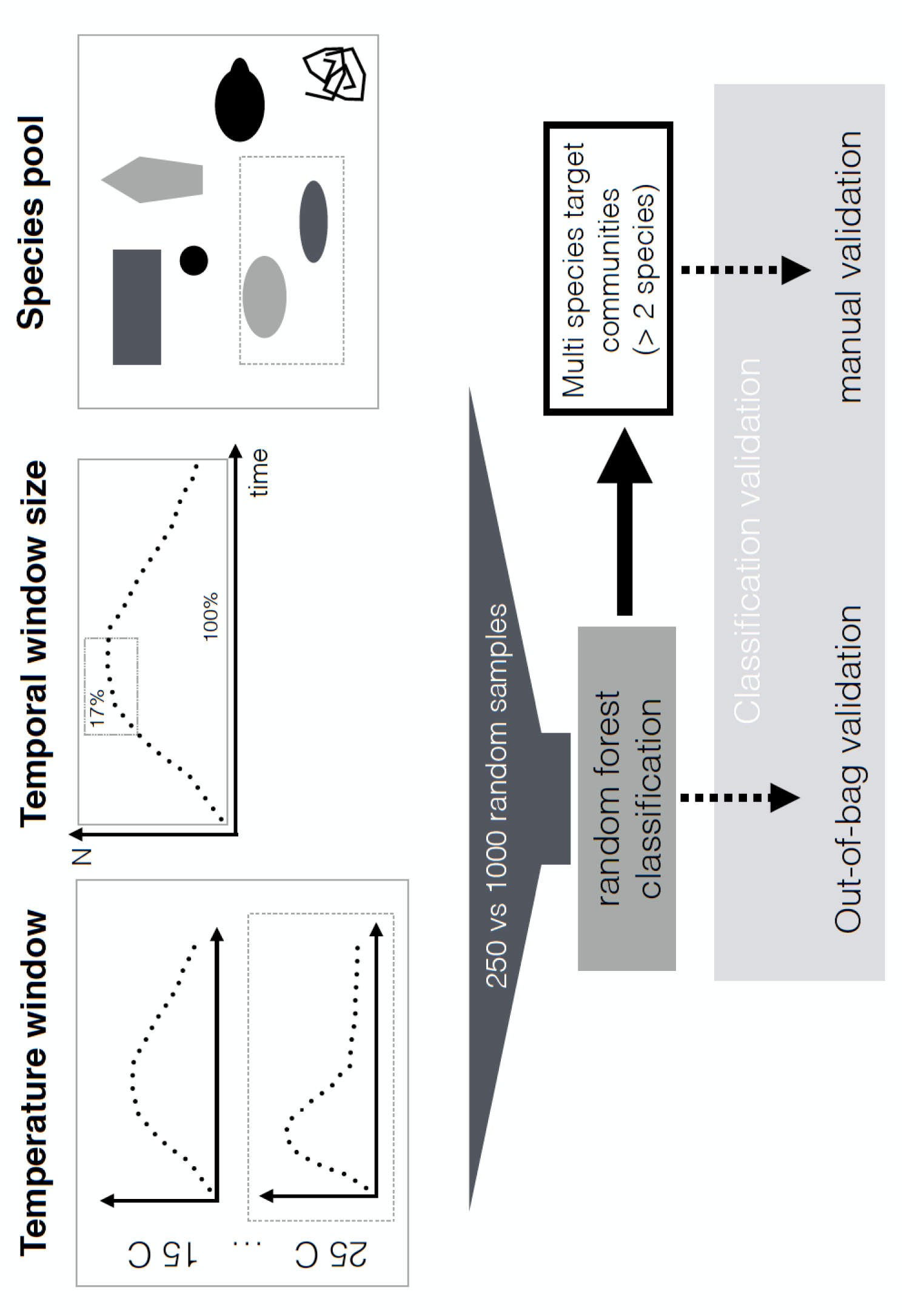
Selecting observations only within certain temperature and temporal distance and using different species pools using the sliding window approach for training the classifier. Imbalance was accounted for via random sub-sampling and the classification results tested using out-of-bag and manual validation.

#### 2.4.6 Fitting the random forest classifier

We used the random forest classifier (RF), as it is computationally efficient and robust, yielding more reliable results than the Naive Bayes and Support Vector Machines, which were also tested (results not presented). Random forest is a widely used classification algorithm based on ensembles of decision trees (Breiman 2001). We used the implementation in the randomForest package in R (Liaw and Wiener 2002). Decision trees are based on binary thresholds that divide the observations into classes, with the goal of the purest possible classes at the end nodes. Features of individuals whose class is known *a priori* are used to train the classifier. The robustness of RF against over-fitting is due to a constrained number of observations and variables used when building individual decision trees, effectively de-correlating trees within the larger ensemble. Each decision tree in the ensemble will predict the class of the unknown event and the final class is based on the majority vote of the ensemble (Cutler et al. 2007).

As there was no association between the replicate of the training data and the test data, we pooled replicates of a given community for training and testing . We grew an ensemble of 500 decision trees for each target community and species identities were assigned to each trajectory according to the majority vote of the ensemble. At each split the RF classifier choose the square root of available variables, and we set the minimum number of observations in each terminal node to one.

### 2.5 Evaluating automatic classification and sliding window approach

We use the out-of-bag classification success in the training data as our response (Breiman 2001). The out-of-bag-success states how well the classification model performs on observations not included in training the model (i.e., a out-of-sample prediction) and hence represents an unbiased measure of classification success. The proportion of individual in the total population correctly predicted as a given species was the response. For the statistical analysis, we used generalized linear mixed models using the lme4 package (Bates et al. 2015) in R (R Development Core Team 2016). Predictor variables were centered and scaled to compare effect sizes and to assist numerical convergence of the models.

First we fitted a model to understand the effect of temperature and species richness on classification success across the full dataset: temperature and richness were modeled as fixed effects; species nested in community composition was included as a random effect to account for the repeated measurements. We also added an individual level random effect (based on the microcosm ID) to account for the over-dispersion in the data (Harrison 2015).

To understand the effect of using sliding windows only including part of the training data, we used contrasts among species pool, temporal and temperature window, as well as number of trajectories randomly sampled. For these contrasts, only classification success from the pairwise interactions was used. Training and classifying all combinations (> 2 species) would haven taken an excessive amount of time (up to a week for each contrast) and hence only using pairwise interactions allowed us to screen the parameter space in a reasonable amount of time. For each of our four window treatments, we fitted a separate model: temperature and sliding window selection were modeled as fixed effects. As before, species nested in community composition was included as a random effect and an individual level random effect (based on the microcosm ID) accounted for excessive over-dispersion in the data (Harrison 2015).

### 2.6 Validation against manual classification

The automatic classification approach was validated against manual identification of species by experts. We randomly selected 3 trajectories for each species, from richness level 3 using samples from days 14, 25 and 37 after the start of the experiment and from three temperatures (15 °C, 21 °C and 25 °C). Three experts independently assigned individuals (i.e. trajectories) to species. We used a majority vote (e.g., majority of votes of the different human observers and the automatic identification) as the reference against which we tested the identifications of each expert. The majority vote was not always unanimous. In 613 of 661 trajectories a majority vote was established, whereas in the remainder no majority vote was found and hence trajectories discarded from further analysis. 37.5% of these cases were IDs divided between *Tetrahymena/Dexiostoma*, *Dexiostoma/Loxocephalus* (14.5%) and *Colpidium/Loxocephalus* (12.5%).

We evaluated the sensitivity and specificity for each species by comparing each voter against the consensus vote using the confusion matrix (Kuhn 2008).

In a two species classification, sensitivity is defined as:

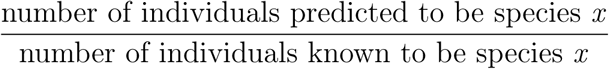

whereas specificity is defined as:

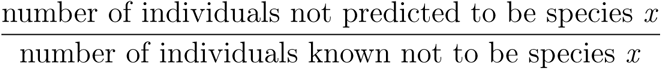

## 3 Results

### 3.1 Video processing and analysis

Of the 18 720 acquired videos, 165 were excluded to due to contamination resulting in a total of 18 555. 18 320 of these were successfully processed to provide particle morphology (species identification) and movement features (Fig S1). The 235 unprocessed videos contained excessive debris (many thousands of particles per frame) compared to the processed videos, causing the particle tracking algorithm to fail. These high particle numbers resulted from directional flow of liquid in the microscope slide generally caused by improper handling or external disturbances during the video recording. The dataset from the processed videos contained 1 702 138 177 observations across 43 267 551 trajectories.

### 3.2 Training data curation

Monocultures were used for training the classifier and hence needed to be of the highest quality. We removed an additional 45 videos from the controls, and 164 from the monocultures to exclude videos with moving background or videoing errors, resulting in a total of 18 111 videos for the final analyses.

The stronger filtering applied to the training data resulted in removal of 3 897 680 out of 4 214 617 trajectories (92% reduction). Most of the removed trajectories were very short and represented random noise associated with floating debris or very short trajectories. The morphological boundaries for each species (see table S2) applied to the remaining 316 937 trajectories removed another 22 230 trajectories from the monoculture data resulting in 294 707 trajectories analysed for training the classifier.

### 3.3 Feature selection and pre-processing

Seven principal components (PCs) accounted for about 95% of the variability in the data. PC1 is strongly associated with the eight features relating to cell size, all having positive associations (Fig 3). PC2 is related to variability in turning angles, size and shape. PC3 is strongly negatively associated with speed features. PC4 captures mostly the mean aspect ratio, whereas PC5 to PC7 are only weakly associated with original features.

### 3.4 Noise identification with Gaussian mixture models (GMMs) and exclusion of background noise

Noise was removed from each subset of data used for training. Trajectories from control cultures (no ciliates), occupied a distinct area of PC1 versus PC2 feature space (S2). The presence of these trajectories in the communities with ciliates sometimes created two relatively distinct clouds of trajectories (e.g., *Paramecium*), and sometimes overlapping clouds of trajectories (e.g., *Dexiostoma*), respectively in Fig S2. Importantly, despite noise distributions overlapping certain species distribution in a subset of dimensions, its location was likely to be different in some of the other dimensions (not plotted).

**Figure 3.**
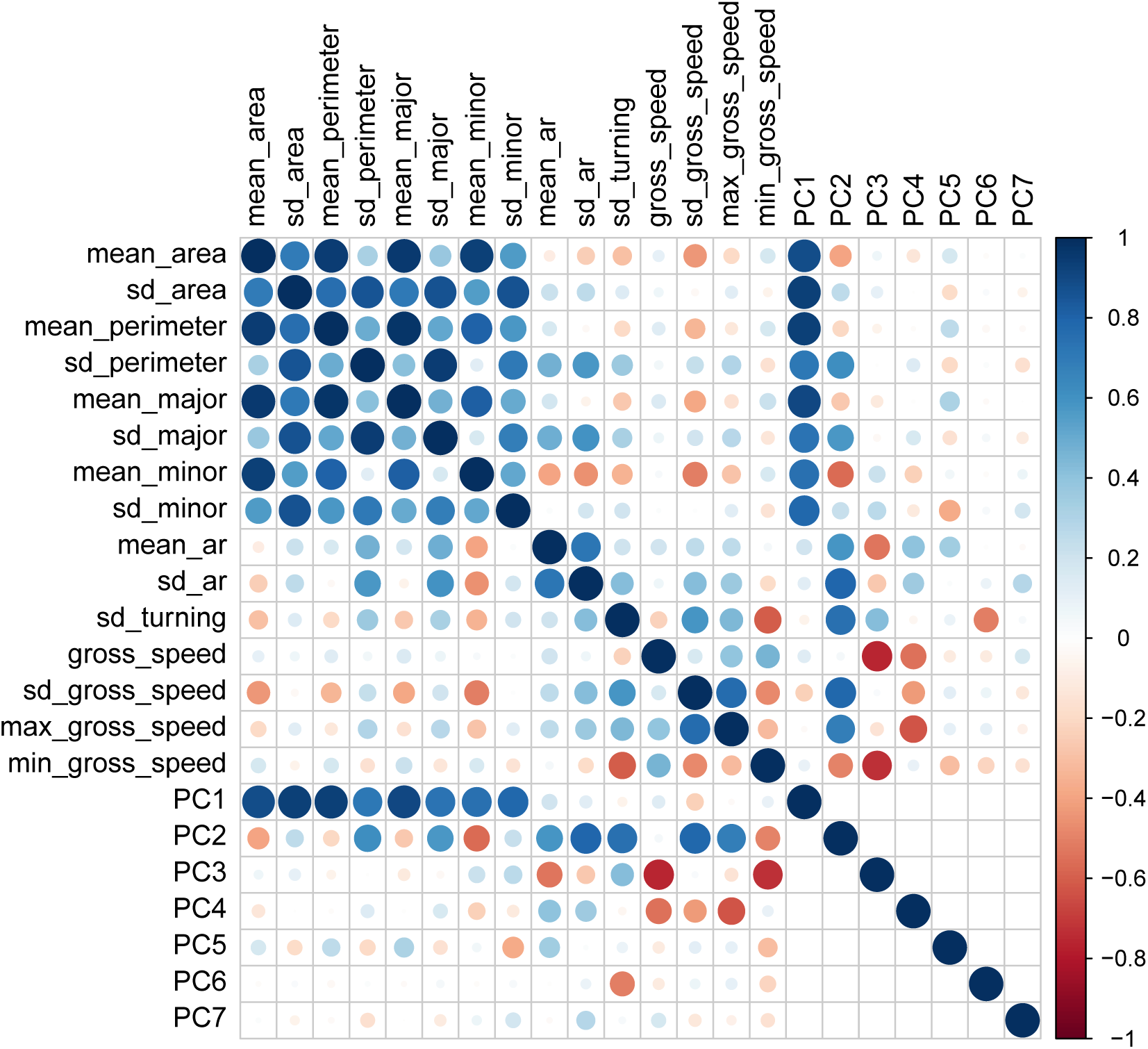
Correlations among original features and principle component scores.

The GMM was able to identify noise and ciliate populations (Fig S3). Trajectories from the training data (“Tetra” or “Loxo”) falling into the area with spurious trajectories from the empty communities (“none”), were re-classified as noise. Reclassification resulted in fewer trajectories from the training data residing in the “noise region” of feature space (Fig S3). The 90% confidence ellipse fitted around the observations helped to remove extreme outliers and improved the species boundaries in multivariate trait space (see Fig S4) .

### 3.5 Effects of temperature, species richness, and sliding window on classification success

Overall, increased temperature (b = -0.129, SE = 0.016, p < 0.001) and species richness (b = -0.852, SE = 0.119, p < 0.001) decreased classification success across species combinations, whereas their interaction was non-significant (b = -0.012, SE = 0.016, p = 0.47; table 3). The richness effect was about seven-fold stronger than the temperature effect (table 3 and Fig 4).

Looking at the contrasts, classification success decreased when all species were included in the training species pool compared to only the known species comprising the community (b = -0.871, SE = 0.003, p < 0.001). Temperature decreased classification further (b = -0.127, SE = 0.023, p < 0.001) and the interaction between temperature and species pool was also negative (b = -0.032, SE = 0.004, p < 0.001; table S3).

Increasing the temporal window size decreased classification (b = -0.080, SE = 0.004, p = 0.001), supporting that smaller temporal windows are beneficial because they capture the temporal dynamics. However, the effect was weaker than the temperature effect (b = - 0.104, SE = = 0.041, p < 0.05) and not further mediated by temperature (b = -0.0003, SE = 0.004, p = 0.93; table S4).

**Table 3.** Model output comparing effect of temperature and species richness on classification success

|  | Model 0 |
| --- | --- |
| (Intercept) | 3.056 (0.119)*** |
| temperature | -0.129 (0.016)*** |
| richness | -0.852 (0.119)*** |
| temperature:richness | -0.012 (0.016) |
| Num. obs. | 24248 |
| Num. groups: ID | 282 |
| Num. groups: combination:predicted.species | 156 |
| Var: ID (Intercept) | 0.063 |
| Var: combination:predicted.species (Intercept) | 2.153 |

\*\*\* $p < 0.001$ , \*\* $p < 0.01$ , \* $p < 0.05$

**Figure 4.**
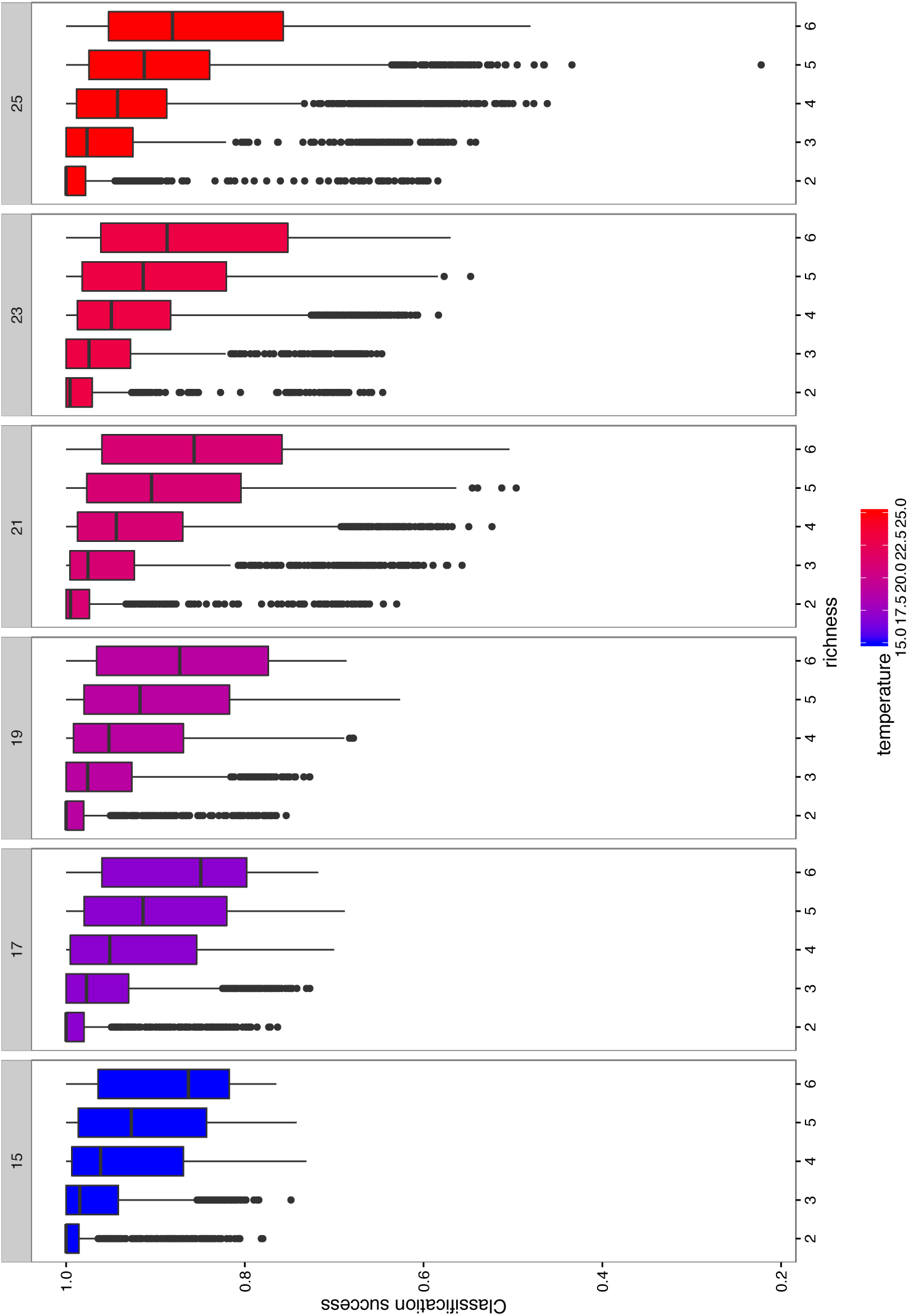
Observed classification success across all the experiment. Both species richness (x axis) and temperature (panel) decreased classification success. At higher temperatures, certain combinations drop in classification success resulting in slightly lower mean classification success remains relatively stable across temperatures.

Including more temperatures in the training data decreased classification success (b = -0.072, SE = 0.004, p < 0.001), and the effect size was similar to the temperature effect itself (b = -0.076, SE = 0.019, p < 0.001). The interaction between temperature and number of included temperatures was positive suggesting that these effects cancel out (b = 0.075, SE = 0.005, p < 0.001; table S5).

Finally, classification success increased with the number of trajectories included (b = 0.141, SE = 0.004, p < 0.001), whereas the temperature effect was negative (b = -0.135, SE = 0.052, p < 0.001). No interaction effect was found meaning that higher numbers of trajectories generally were beneficial across temperatures (b = 0.006, SE = 0.004, p = 0.11; table S6).

### 3.6 Validation against manual classification

When we compared the classification success for automatic and manual classification, we observed high classification success (sensitivity and specificity) for both manual and automatic classification of the six ciliate species (Fig 5). Manual classification is often slightly better than automatic classification, but automatic classification can outperform manual observers for some classes (e.g., *Tetrahymena*) (Fig 5). Although we included trajectories from different combinations and temperatures, species classification success remained above or close to 80%, even for species like *Tetrahymena* whose accuracy was lower in the out-of-bag validation. Furthermore, the data suggests that sensitivity is correlated between manual and automatic classification, i.e. that they experience the difficulties with the same species.

Automatic classification did less well in identifying spurious trajectories (i.e., noise), with *Tetrahymena* (39%) and *Paramecium* (11%) being the most confounded classes. Although this shows that some noise escapes our cleaning procedure and that the error is non-randomly distributed across species, overall we found a strong positive correlation between automatic and manual counts of ciliates (Pearson correlation coefficient = 0.86).

## 4 Discussion

Here we introduce a methodological framework to automate species identification from individual phenotypes in dynamic contexts. We show that we can reliably classify species from video recordings using interspecific phenotypic variation in morphology and movement, while accounting for environment dependent intraspecific phenotypic variation. We developed and optimized our approach on identifying ciliates in aquatic microcosm communities, but the techniques generalize to a wide range of study systems, including aquatic algae and zooplankton. Our results highlight the importance of the dynamic context of biotic interactions and abiotic environment for accurate species level classification. Overall, increased temperature and species richness reduced classification success, with species richness inducing an order of magnitude greater decrease than temperature. Importantly, sub-setting training data according to ranges of time, temperature and species richness yielded increased classification success and mitigated the problem of imbalance.

**Figure 5.**
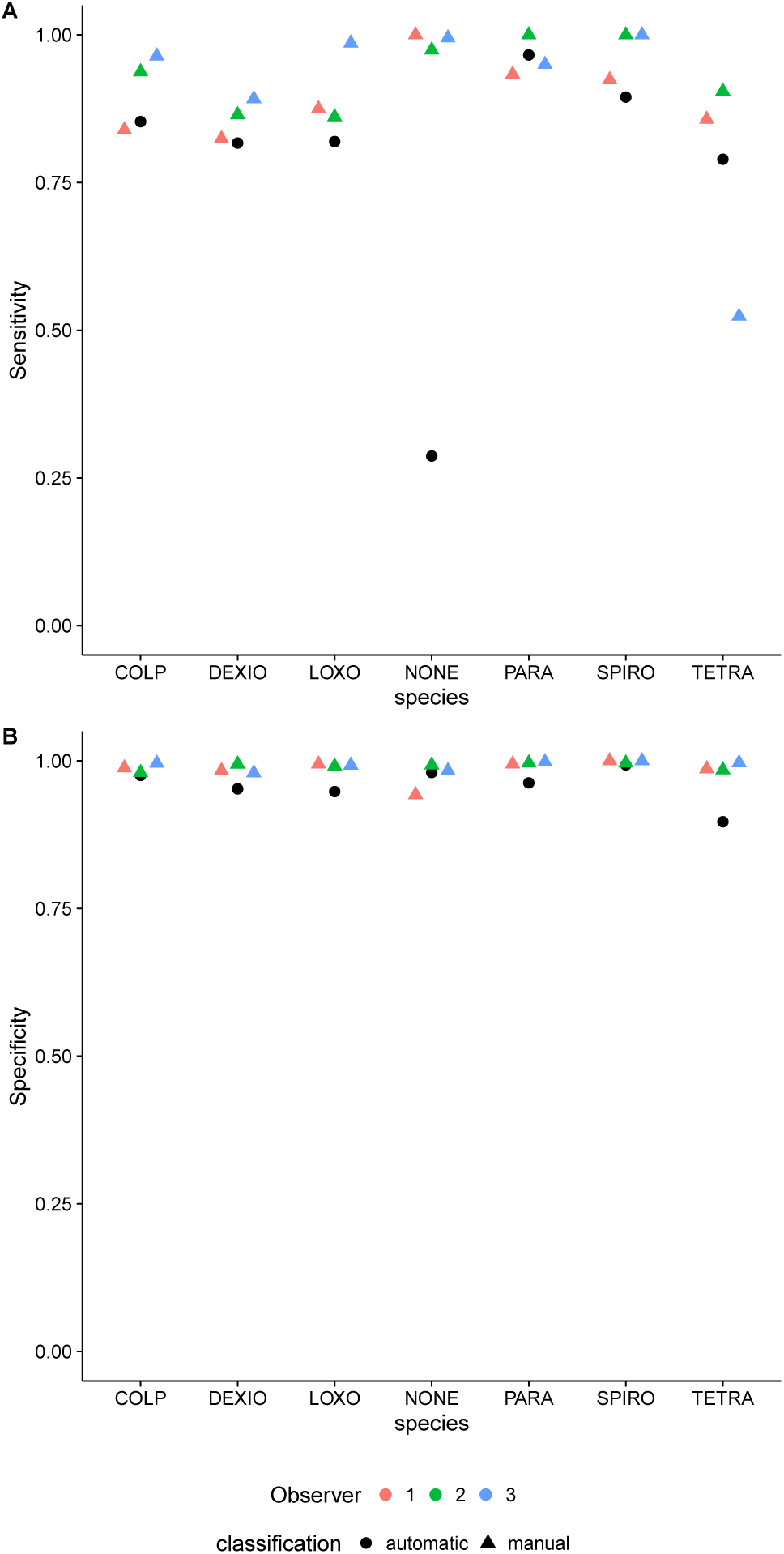
Comparison of manual and automatic classification success for each of the species. Panel A shows the sensitivity, whereas Panel B shows the specificity against the consensus vote. Different colours show different experts, whereas different shapes show manual versus automatic identifications. The automatic classification behaves very similar to the experts both in terms of sensitivity and specificity for the six ciliate species.

### 4.1 The need to account for dynamic trait change in communities

Intraspecific variation in phenotypic expression is routinely observed across the kingdom of life and is attributed to a wide range of abiotic and biotic environmental factors (Ozgul et al. 2009). Specifically, phenotypic plasticity is the mechanism organisms use to cope with environmental variation that allows for changes in morphology, physiology or behavior and may encompass acclimation or epigenetic responses (Price et al. 2003). We used ciliates as study organisms, as they show large variation in phenotypic response to environmental factors, for instance smaller body size to increased temperature (Atkinson et al. 2003) or changes in movement/feeding behavior to the presence of predators/competition (Kusch 1993). Despite these changing characteristics over time and space, we show that our automated pipeline can accurately identify species across considerable ecological complexity. This highlights the importance of accounting for the range of phenotypes when identifying species based on morphology (Vrijenhoek 2009).

### 4.2 Improved trait resolution due to dynamic sliding windows

The dynamic sliding window approach was superior to models that include global information (e.g., using all species, all dates, or temperatures for training purposes), when classifying multi-species communities. The smallest temporal window showed higher classification success compared to using all dates to determine classes. Additionally, limiting the classification to specific species and temperatures improved classification. Improved classification performance is due to the ability of our framework to capture changes in traits (i.e. phenotypes) due to the environment (e.g. species interactions and temperature) in the training subsets. In contrast, if data were pooled without considering the environment, systematic changes would be swamped by the overall trait variability.

Our finding can be easily illustrated with a trait like body size. Imagine two species that initially differ in mean body size, that shrink through time, but at different rates (i.e. resulting in overlapping body sizes). If all data were pooled, the classifier would consider individuals within the whole range of body sizes observed over the experiment for the two species, due to the overlapping body sizes. However, a classifier that only considers body size at a given time point exploits temporally non-overlapping sizes more efficiently and may hence have higher classification success.

### 4.3 Community context and potential to tell apart similar species

The model including only the expected species in a community performed far better than the model including all species. This is expected, as classifiers tend to perform less well with increasing numbers of classes to predict (Kuhn and Johnson 2013). The corollary is that if the presence of certain species can be ruled out (as in our experimental design), it is better to work with a more specific classifier. This may not work under non-experimental settings where the expected species are unknown and also has costs in terms of re-training the classifier multiple times. If the species to be expected in a community are unknown, training the classifier on all possible species may be a better choice.

Our results highlighted difficulties in telling the two smallest species apart ( *T*. *thermophila* and *D*. *campylum*). This effect was pronounced at higher temperatures, potentially due to decreased size at higher temperatures (Atkinson et al. 2003). A higher magnification for communities containing the smallest ciliates has potential to improve classification as individual traits such as cell shape may be better resolved. However, higher magnifications entail a decrease in the volume sampled and hence this trade-off needs to be carefully balanced.

Another avenue for telling phenotypically similar species apart could be to apply active learning approaches to the training of the classifier (Sommer and Gerlich 2013). Active learning can improve the boundaries between classes by using user input on decisive observations. Instead of just increasing the overall amount of observations available, this technique identifies observations that are critical in assigning a large number of other observations to a given class. The active learning algorithm selects observations autonomously and presents them to the human expert for annotating (Jones et al. 2009). The manual validation has shown that experts can provide reliable identifications from videos, and the costs of manual identifications may be paid off by substantial gains in classification success when decisive training observations would be manually confirmed.

### 4.4 Data curation, cleaning, feature selection and dimension reduction

Much of the data cleaning involved careful validation of the raw data, identifying potential problems with the data and designing steps to subsequently clean the data in a more automated fashion. Fundamentally, the classifier is only as good as the training data, meaning that foremost the quality and then the quantity is important. Observations accidentally labeled as another class (e.g., spurious trajectories due to moving background confounded with ciliate individuals) may seriously hamper the classification. Our cleaning pipeline therefore deliberately discarded a large amount of trajectories (> 92%) in the first step. This amount is nevertheless comparable to other automatic classification pipelines, for instance, for marine planktonic organisms, where more than 95% of particles were discarded (Bi et al. 2015) before target objects are classified.

Reducing the number of features had the advantage of requiring less computing time when training the classifier, especially when predictors are highly correlated and hence contain the same information. An excessive number of features may have also decreased the accuracy of the classification, a phenomenon known as the curse of dimensionality (Sommer and Gerlich 2013). Albeit RF classification does not require feature reduction and transformations to work, better numerical stability is expected when features are on the same scale (Kuhn and Johnson 2013).

### 4.5 Down-sampling data for training

Randomly sampling a number of trajectories from the subsets (training data) reduces the chance of over-fitting as well as removes bias from the random forest classifier because all classes have approximately the same number of training cases. This proved to be a problem in previous applications of RF (Pennekamp et al. 2015, Soleymani et al. 2015), where minority classes often had lower classification success than majority classes. Whereas some classifiers are more robust to imbalance (e.g., support vector machines), here we show that sub-sampling the population circumvents the imbalance issue. However, sufficient observations of the minority class are still needed. Increasing the number of observations increased classification success, though with computational cost. Training the GMM on 1000 instead of 250 trajectories led to a four-fold longer training time in the two species combinations, and potentially much longer training in more species rich communities. In case the observations of the minority class are limiting, it may also require increasing the temporal window with associated decreases in classification. Balancing these factors hence needs careful consideration of the properties of the classification problem studied.

### 4.6 Speed, size of experiment, scalability, workflow

Video analysis and automated species identification allowed a much larger experiment than could have been achieved with manual methods, with collection of much more (and different data). Manual identification took 3 to 4 person hours for the limited number of trajectories (<700) in the validation dataset, and hence manual classification of all the trajectories in the dataset would have been impossible (>40 000 000 trajectories). Manual counting of the samples under the microscope can take 5-10 minutes depending on abundance and species richness, requiring up to 120hours of continuous work on each sampling date; this is not feasible. This high person-hour demand associated with extensive multispecies microcosm experiments limits sampling events to either time series data for one ciliate species across a limited number of microcosms (50 to 200) (e.g. Leary and Petchey 2009, Jiang and Morin 2004, Petchey et al. 1999, Seymour et al. 2015), or more experimental units (>300) but few sampling occasions (e.g. Carrara et al. 2015b;a). Our approach circumvents some of these logistic limitations and furthermore gathers rich phenotypic information that opens microcosm experiments, and other similar study systems, to high-throughput analyses of traits.

### 4.7 Caveats and limitations

Out-of-bag error rates showed high classification success for ciliate and background noise. The manual classification showed that automatic classification is indeed comparable to manual classification for the six ciliates. However, a certain amount of trajectories identified as noise by the manual observers were classified as ciliates by the RF. A possible explanation is that training data was deliberately processed to contain the most reliable observations of a given class. The strong filtering removed most of the noise, but also ciliate trajectories, probably improving the separation of classes in multivariate trait space. In the test data, no strong filtering was applied to make sure all ciliate trajectories remained, however, this also led to spurious trajectories being identified as ciliates. Whereas we miss out on some noise, bias in the species counts should be negligible for several reasons. First, the strong correlation between observed and predicted counts based on the validation dataset indicates that the automatic classification provides reliable counts. Second, the counts in a given community at a given date are based on a weighted average: a given identification contributes only to the number of frames it was detected on. Most of the trajectories that were wrongly classified by the RF, were substantially shorter than the correct identifications and hence can only contribute a small fraction to the overall count. Third, the quantification of different error sources allows us to incorporate specific measures of observational error into models to analyze the data in a later step. Our manual validation also showed that human observers do not agree on identifications unanimously, however, the human error is almost never stated nor quantified and hence cannot be considered in subsequent analyses.

## 5 Conclusions

Our analysis framework based on sliding windows allows reliable classification of individual organisms into species, despite temporal and environmentally induced trait change. We developed the approach based on videos of ciliate species, but the methodology and computational pipelines are general and hence applicable to a wide range of organisms, for example to monitor algae communities and the dynamics of microbial organisms in biofuel production (Pons and Vivier 2000), track diversity in freshwater or marine plankton (Biard et al. 2016), or in sewage plants (Amaral et al. 2004; 2008).

## Acknowledgments

The data used in this study was collected in a joint effort of the Altermatt and Petchey groups, University of Zurich.

We thank S3IT at University of Zurich for adapting the tracking functionality of the R package BEMOVI to the ScienceCloud computing infrastructure and assistance in processing the video raw data.

FP and OLP were financially supported by Swiss National Science Foundation Grant 31003A_159498, FA and MS received funding by the Swiss National Science Foundation Grants 31003A_135622 and PP00P3_150698.

